# SMURF2 inhibits autophagic control of *Mycobacterium tuberculosis* in macrophages

**DOI:** 10.1101/2025.09.02.673599

**Authors:** Priscila C. Campos, Kathryn C. Rahlwes, Victoria A. Eknitphong, Beatriz R.S. Dias, Kubra F. Naqvi, Samuel Alvarez-Arguedas, Michael U. Shiloh

## Abstract

Autophagy is a critical host defense mechanism that restricts intracellular pathogens such as *Mycobacterium tuberculosis* (Mtb). A key step in this process is the ubiquitination of Mtb or Mtb-associated structures. The E3 ligase SMURF1 catalyzes K48-linked ubiquitination, promoting bacterial clearance. However, the function of its homolog, SMURF2, in host defense remains undefined. Here, we demonstrate that *Smurf2* deletion in murine macrophages increases SMURF1 levels, enhances LC3B lipidation, augments K48 ubiquitination of Mtb-associated structures, and reduces intracellular Mtb replication. These effects are reversed by *Smurf1* deletion, indicating that SMURF2 restricts autophagy in a SMURF1-dependent manner. Mice with myeloid-specific *Smurf2* deletion exhibit modestly prolonged survival following aerosol Mtb infection. In human macrophages, *SMURF2* knockdown or its pharmacological inhibition with the HECT ligase inhibitor Heclin reduces Mtb replication. Together, our findings identify SMURF2 as a negative regulator of selective autophagy and host immunity to Mtb and suggest that targeting SMURF2 may represent a novel host-directed therapeutic strategy for tuberculosis.

## Introduction

*Mycobacterium tuberculosis* (Mtb), the causative agent of tuberculosis (TB), is responsible for more than 10 million new cases and 1.3 million deaths each year, making it the leading cause of death from a single infectious agent worldwide (1). The airborne route of transmission, long latency period, and capacity for immune evasion all contribute to the persistence and global impact of TB. A recent analysis projected over 31 million TB-related deaths between 2020 and 2050, with a corresponding global economic loss of 17.5 trillion US dollars (1).

The standard treatment for TB relies on prolonged multidrug chemotherapy, typically requiring at least six months of adherence for drug-susceptible strains. This extended treatment duration, combined with limited diagnostic tools for early detection of drug resistance, has contributed to the emergence of multidrug-resistant (MDR) and extensively drug-resistant (XDR) Mtb strains (2). These challenges underscore the need for innovative therapeutic strategies that not only shorten treatment but also enhance host immune control of infection. Host-directed therapies (HDTs) aim to bolster the host immune response rather than directly targeting the pathogen, offering the potential to synergize with antibiotics, improve treatment outcomes, and overcome drug resistance (3, 4). However, realizing the promise of HDT requires a more detailed understanding of host cell-autonomous mechanisms that limit Mtb replication.

Autophagy is a conserved intracellular degradation pathway that maintains cellular homeostasis and promotes clearance of intracellular pathogens (5, 6). When the autophagic machinery encounters intracellular pathogens, the ubiquitin-proteasome system (UPS) is engaged through the attachment of ubiquitin (Ub) molecules onto intracellular cargo, targeting the cargo for degradation (7–10). Ubiquitinated intracellular cargo is recognized by cargo receptors, such as sequestrosome-1 (SQSTM1/p62), calcium-binding and coiled-coil domain-containing protein 2 (CALCOCO2/NDP52), optineurin (OPTN), and neighbor of BRCA1 gene 1 (NBR1). These receptors contain both ubiquitin-binding domains (UBDs) and LC3-interacting regions (LIRs) that enable cargo delivery to LC3-decorated autophagosomes for subsequent degradation following lysosomal fusion (10–13).

The ubiquitination cascade is mediated by the sequential action of E1 activating enzymes, E2 conjugating enzymes, and E3 ligases, with E3 ligases conferring substrate specificity (14). A limited number of E3 ligases have been identified that ubiquitinate Mtb-containing structures with important roles in xenophagic control of infection (15–17). For example, tripartite motif-containing (TRIM) E3 ligases such as TRIM32 enhances autophagy in Mtb-infected human macrophages (18), PARKIN mediates K63-linked polyubiquitination and recruitment of p62 and NDP52 to Mtb-containing structures (19) and Smad-specific E3 ubiquitin protein ligase 1 (SMURF1) promotes K48-linked ubiquitination and proteasome recruitment (10), highlighting distinct and non-redundant mechanisms of bacterial clearance (20).

E3 ligases are regulated by post-translational modifications such as phosphorylation, acetylation, sumoylation, and ubiquitination (21, 22). In breast cancer cells, SMURF2 negatively regulates SMURF1 by inducing its ubiquitination and degradation (23). SMURF1 and SMURF2 share a conserved structure with an N-terminal C2 domain for membrane binding, WW domains for protein-protein interactions, and a C-terminal HECT (homologous to the E6-AP carboxyl terminus) ubiquitin-ligase domain (24). Initially identified as negative regulators of the transforming growth factor-β (TGF-β) and bone morphogenetic protein (BMP) signaling (25, 26), SMURF1 and SMURF2 have since been implicated in diverse biological processes, including bone metabolism and host-pathogen interactions (10, 27, 28).

Although SMURF1 and SMURF2 share structural similarities and partially overlapping substrates, they also exhibit distinct functions. SMURF2 can induce the degradation of SMURF1, whereas SMURF1 does not affect SMURF2 stability (23). While both ligases are well studied in cancer biology (29, 30), their roles in cell-autonomous immunity remain poorly understood. Here, we demonstrate that genetic deletion of *Smurf2* in murine macrophages significantly enhances SMURF1 levels, LC3B lipidation, and autophagic targeting of Mtb, resulting in reduced bacterial replication. We further show that myeloid-specific *Smurf2* deletion provides modest protection in a murine TB model and that SMURF2 inhibition in human macrophages, both genetically and pharmacologically, enhances bacterial clearance. These findings identify SMURF2 as a negative regulator of cell-autonomous immunity to Mtb and a potential target for host-directed therapy in tuberculosis.

## Results

### *Smurf2* deletion in BV2 macrophages increases SMURF1 and LC3B-II levels and reduces intracellular Mtb growth

To examine the role of SMURF2 in cell-autonomous immunity, we used CRISPR/Cas9 to knock out *Smurf2* in the BV2 mouse microglial cell line. We selected BV2 cells for their suitability in genetic manipulation and their prior use in studies of autophagy and mycobacterial pathogenesis (31–38). SMURF2-deficient BV2 cells showed a 60% increase in SMURF1 protein levels compared to wild-type controls **(Figure 1A, 1B)**. To determine whether SMURF2 regulates intracellular Mtb replication, we infected BV2 wild-type and *Smurf2* knockout (KO) cells with Mtb and measured bacterial burden by colony-forming unit (CFU) assay. While Mtb grew steadily in control BV2 cells, BV2 *Smurf2* KO cells exhibited significantly reduced bacterial growth at days 2 and 3 post-infection **(Figure 1C).**

**Figure 1:**
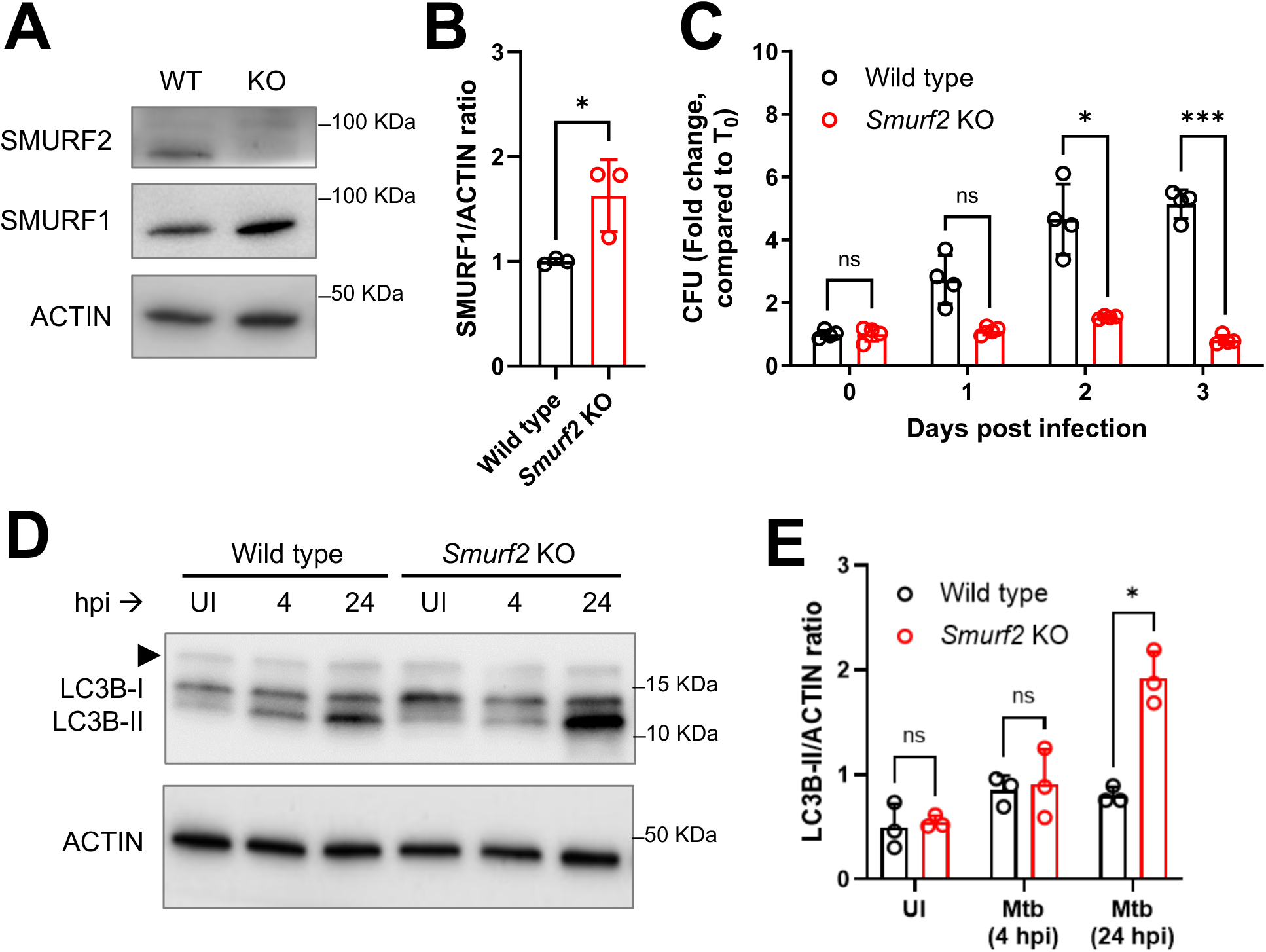
Murine macrophages lacking *Smurf2* are more resistant to Mtb and show an increased LC3B lipidation. A) Representative immunoblot analysis of SMURF2 and SMURF1 accumulation in BV2 wild type (WT) and BV2 *Smurf2* knockout (KO) cells under basal conditions. B) Densitometric quantification of SMURF1 levels, as shown in (A) (mean ± SD of 3 independent experiments, *p ≤ 0.05; unpaired t-test). C) Intracellular Mtb colony forming units (CFU) in BV2 WT and *Smurf2* KO cells at different timepoints post-infection (mean ± SD of a representative experiment performed in quadruplicate, ns, non-significant, p > 0.05, *p ≤ 0.05 and ***p ≤ 0.001; two-way ANOVA). D) Representative immunoblot of LC3B-II levels in BV2 WT and *Smurf2* KO cells under basal (UI) and at 4- and 24-hours post-infection (hpi). E) Densitometric quantification of LC3B-II levels, as shown in (D) (mean ± SD of 3 independent experiments, ns, non-significant, p > 0.05, *p ≤ 0.05; two-way ANOVA).

We next examined whether loss of SMURF2 affects autophagy. Lipidation of Atg8 family proteins, including LC3B, is a key step in autophagosome formation (39) and is commonly assessed by monitoring the conversion of LC3B-I to LC3B-II (40, 41). LC3B-II levels were quantified by immunoblot in both BV2 wild-type and BV2 *Smurf2* KO cells under both basal conditions and after 4 and 24 hours of Mtb infection. While there were no differences after 4 hours, LC3B-II was increased two-fold in SMURF2-deficient cells at 24 hours after infection **(Figures 1D, 1E)**. These findings suggest that SMURF2 negatively regulates autophagy in Mtb-infected macrophages, potentially through its effect on SMURF1 stability.

### K48-linked ubiquitin colocalization with Mtb is increased in BV2 *Smurf2* KO macrophages

SMURF1 promotes K48-linked ubiquitination of substrates, targeting them for proteasomal degradation. We previously showed that loss of SMURF1 reduces K48-linked ubiquitin colocalization with Mtb in mouse bone marrow-derived macrophages (BMDMs) (10). Since SMURF1 levels were elevated in BV2 *Smurf2* KO cells (**Figure 1A, 1B**), we hypothesized that K48-linked ubiquitination of Mtb-associated structures would be correspondingly increased in these cells. To test this, we infected BV2 wild-type and BV2 *Smurf2* KO cells with Mtb expressing mCherry (Mtb-mCherry) and performed immunofluorescence microscopy to assess colocalization with K48-linked ubiquitin. BV2 *Smurf2* KO cells exhibited a significant increase in K48-linked ubiquitin colocalization with Mtb-associated structures compared to wild-type controls **(Figures 2A, 2B)**.

**Figure 2:**
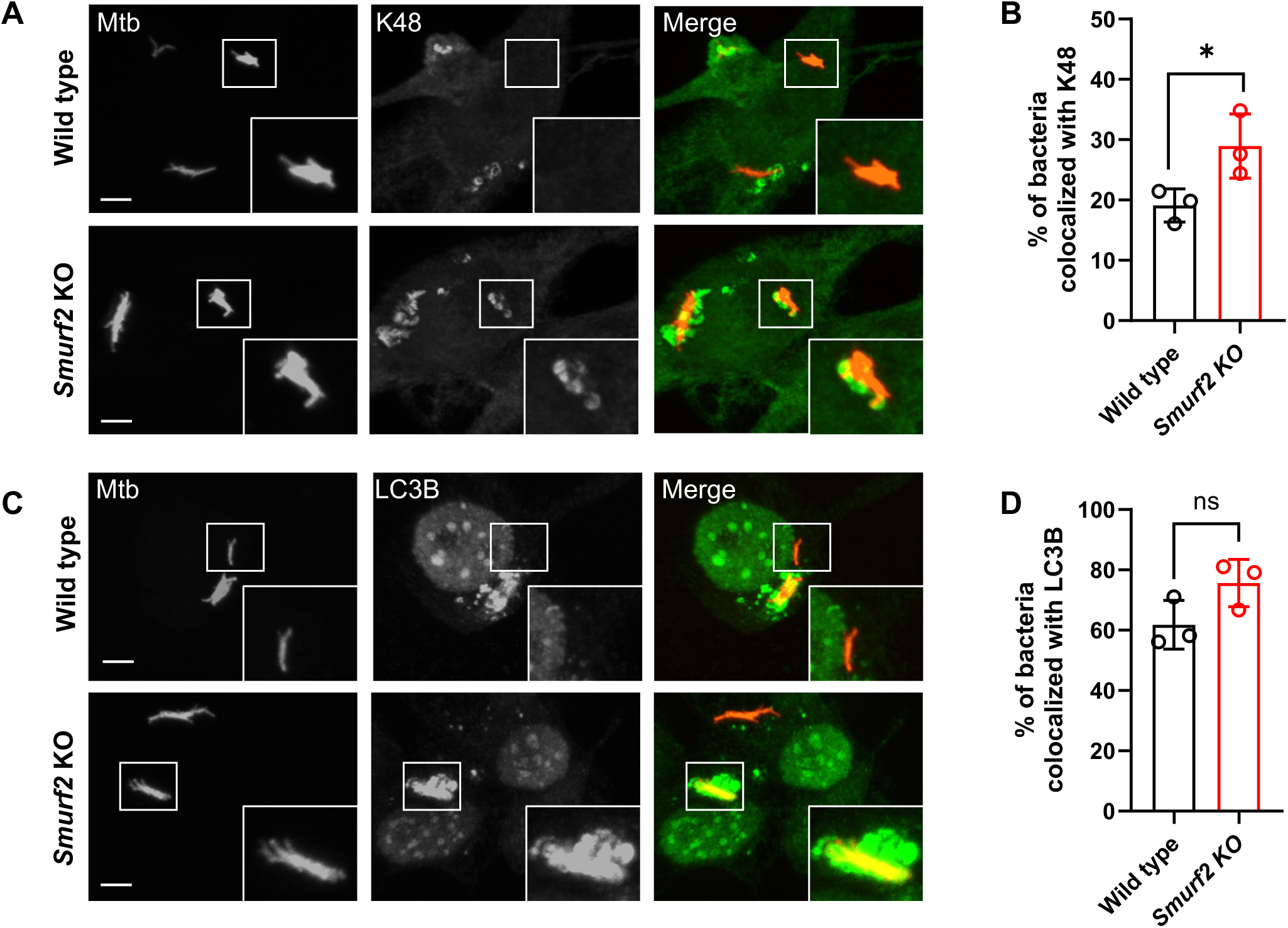
K48 and LC3B colocalize with Mtb in murine macrophages lacking *Smurf2*. A) Representative immunofluorescence images of Mtb-mCherry (red) and K48 ubiquitin (green) in BV2 WT and BV2 *Smurf2* KO cells. Images were captured 17 hours post-infection using an anti-K48 antibody. B) Quantification of Mtb-mCherry colocalization with K48 from (A) (mean ± SD of a 3 independent experiments, performed in triplicate, *p ≤ 0.05; unpaired t-test). C) Representative immunofluorescence images of Mtb-mCherry (red) with LC3B (green) in BV2 WT and *Smurf2* KO cells. Images were acquired 17 hours post-infection using an anti-LC3B antibody. D) Quantification of Mtb-mCherry colocalization with LC3B from (C). (mean ± SD of a 3 independent experiments, performed in triplicate, ns, non-significant; unpaired t-test). Scale bar, 5 µm.

We next evaluated LC3B recruitment to Mtb-associated structures under the same conditions. Although LC3B colocalization was modestly increased in BV2 *Smurf2* KO cells compared to BV2 wild type cells, the difference did not reach statistical significance (**Figures 2C, 2D)**. These data indicate that *Smurf2* deficiency enhances K48-linked ubiquitination of Mtb-associated structures, consistent with increased SMURF1 expression in BV2 *Smurf2* KO macrophages.

### *Smurf2* deficiency impacts Mtb replication in a *Smurf1*-dependent manner

Given that *Smurf2* deletion increased SMURF1 levels and that SMURF1 regulates selective autophagy during Mtb infection (10), we tested whether the phenotype observed in BV2 *Smurf2* KO cells was dependent on SMURF1. We used CRISPR/Cas9 to generate BV2 *Smurf1* KO and BV2 *Smurf2/1* double KO cells and quantified SMURF1 and SMURF2 protein levels at baseline. As expected, SMURF1 was increased in BV2 *Smurf2* KO cells, while SMURF2 levels were unchanged in BV2 *Smurf1* KO cells **(Figure 3A)**.

**Figure 3:**
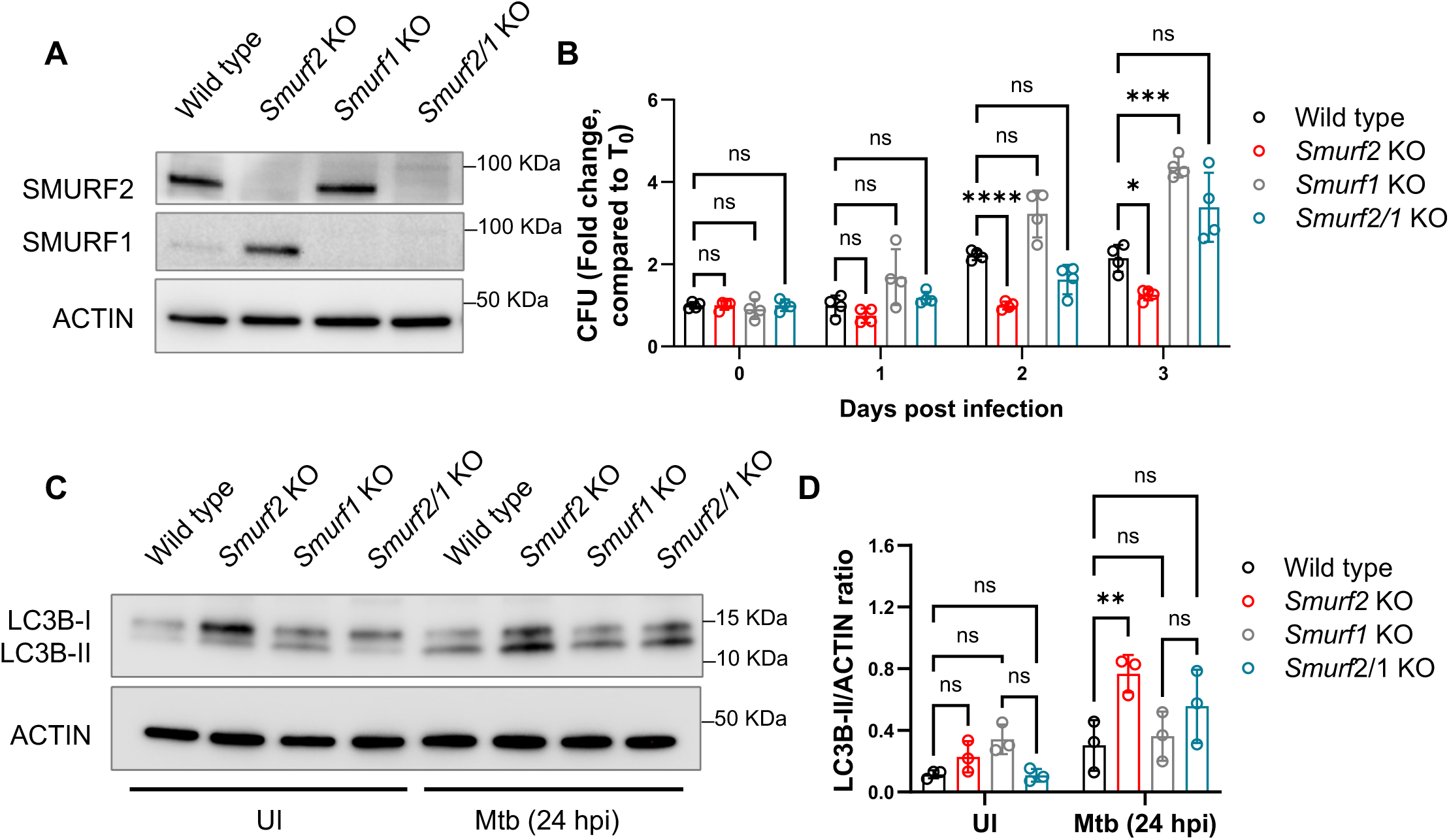
*Smurf2* knockout impacts Mtb replication through SMURF1 in murine macrophages. A) Immunoblot analysis to detect SMURF2 and SMURF1 in BV2 WT, *Smurf2* KO, *Smurf1* KO, and *Smurf2/1* KO cells under basal conditions. B) Colony forming units (CFU) of Mtb in BV2 WT, *Smurf2* KO, *Smurf1* KO, and *Smurf2/1* KO BV2 cells at different timepoints (mean ± SD of a representative experiment performed in quadruplicate, ns, non-significant, *p ≤ 0.05 and ***p ≤ 0.001; two-way ANOVA, with multiple comparisons). C) Immunoblot analysis of LC3B lipidation in BV2 WT, *Smurf2* KO, *Smurf1* KO, and *Smurf2/1* KO cells, infected or not with Mtb (MOI 5) at 24 hours post infection. D) Densitometric quantification of LC3B-II levels, as shown in (C) (mean ± SD of 3 independent experiments, ns, non-significant, **p ≤ 0.01; two-way ANOVA, with multiple comparisons).

We next infected BV2 wild-type, BV2 *Smurf2* KO, BV2 *Smurf1* KO, and BV2 *Smurf2/1* KO cells with Mtb and measured intracellular bacterial burden over three days. Consistent with prior findings in mouse BMDMs (10), BV2 *Smurf1* KO cells showed increased Mtb replication compared to wild-type cells. BV2 *Smurf2/1* KO cells had bacterial burdens comparable to BV2 *Smurf1* KO cells, indicating that the reduction in Mtb growth observed in BV2 *Smurf2* KO cells requires SMURF1 **(Figure 3B)**.

We also examined LC3B lipidation in these cell lines at baseline and after infection. BV2 *Smurf2* KO cells showed increased LC3B-II levels, whereas no change in LC3B lipidation was observed in BV2 *Smurf1* KO or BV2 *Smurf2/1* KO cells **(Figures 3C, 3D)**. This suggests that the effect of *Smurf2* deletion on LC3B lipidation is not mediated by SMURF1. Together, these results indicate that SMURF2 restricts Mtb control through SMURF1-dependent mechanisms and modulates LC3B lipidation through a SMURF1-independent pathway.

### SMURF2 complementation restores susceptibility to Mtb by decreasing SMURF1 protein levels

SMURF2 contains a conserved catalytic cysteine (C716) within its C-terminal HECT ubiquitin ligase domain, which is essential for its enzymatic activity (**Figure 4A**) (42). To confirm that the phenotype observed in BV2 *Smurf2* KO cells was due to loss of SMURF2, we reconstituted SMURF2 expression by lentiviral transduction. In addition, to determine whether SMURF2 suppresses macrophage immunity to Mtb through its ligase function, we generated BV2 *Smurf2* KO macrophages stably complemented with either wild type (WT) SMURF2 (*Smurf2^WT^*) or a catalytically inactive mutant, C716A (*Smurf2^C716A^*), which targets the active site cysteine. Immunoblot analysis confirmed successful re-expression of SMURF2 in the *Smurf2^WT^*-complemented cells. While the catalytic inactive SMURF2^C716A^ mutant was also detected, its band intensity was significantly reduced compared to the SMURF2^WT^, suggesting lower protein abundance (**Figure 4B**). We previously observed that SMURF1 expression is increased in the absence of SMURF2 in BV2 cells (**Figures 1A, 3A**). Consistent with this, complementation with SMURF2^WT^ in BV2 *Smurf2* KO cells reduced SMURF1 levels (**Figure 4C**). In contrast, the SMURF2^C716A^ mutant failed to reduce SMURF1 protein levels, similarly to what was observed in *Smurf2* KO cells with the empty vector (EV) (**Figure 4C**). However, the inability to restore SMURF1 expression in SMURF2^C716A^ mutant expressing cells may also be due to its reduced expression.

**Figure 4:**
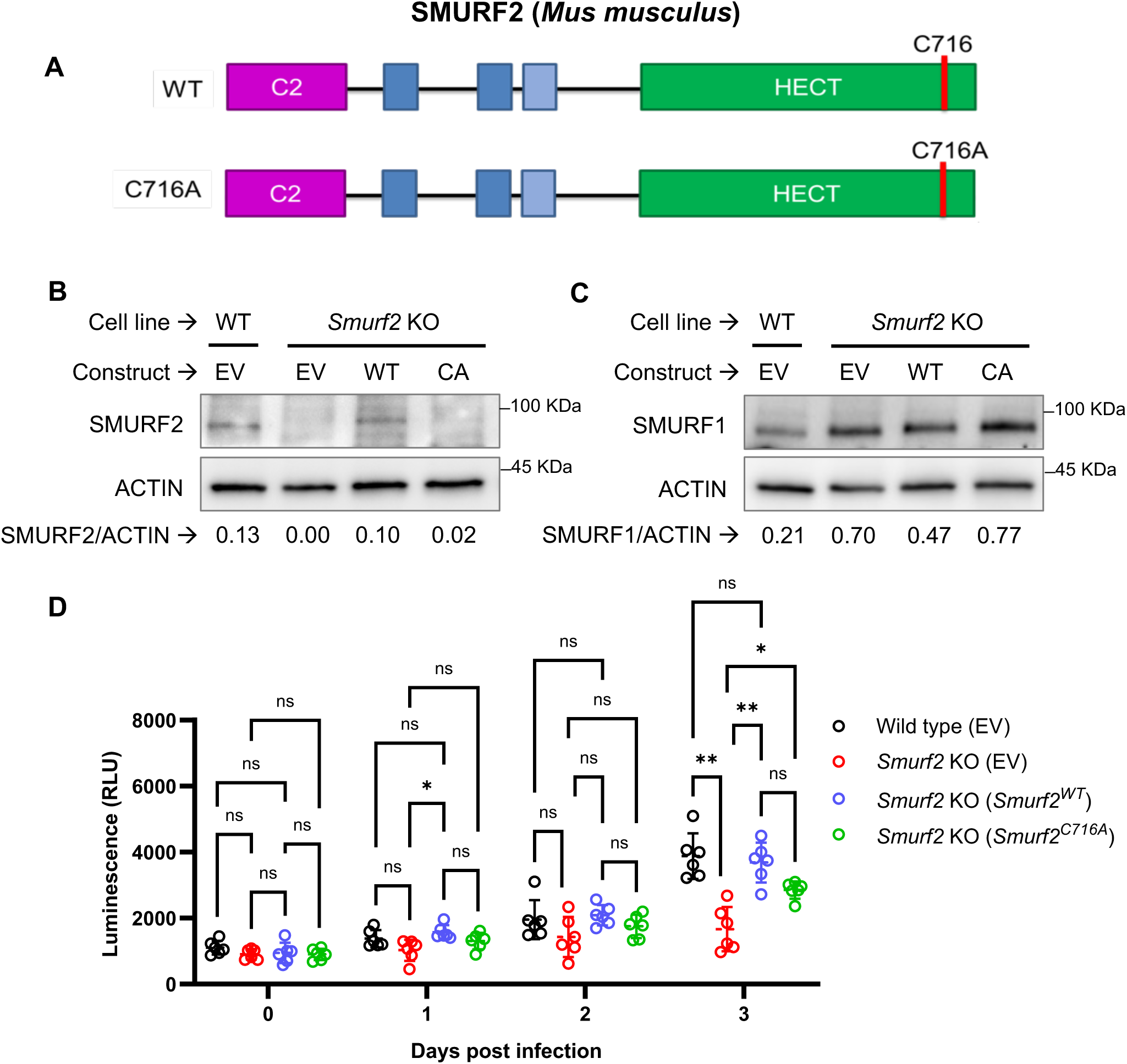
Complementation of BV2 *Smurf2* KO cells for SMURF2 expression using a lentiviral plasmid and impact on Mtb luminescence. A) Schematic of *Smurf2* cDNA constructs cloned into a lentiviral vector, for CMV-driven expression of wild type (SMURF2^WT^) or mutant (SMURF2^C716A^) proteins in BV2 cells. B) Immunoblot analysis of SMURF2 in BV2 cell lysates transduced with either empty vector (EV) or the *Smurf2* constructs shown in (A), under basal conditions, using ACTIN as a loading control with densitometry analysis of SMURF2/ACTIN ratio. C) Immunoblot analysis of SMURF1 in BV2 cell lysates transduced with EV or the *Smurf2* constructs shown in (A), under basal conditions, using ACTIN as a loading control with densitometry analysis of SMURF1/ACTIN ratio. D) Luminescence from Mtb-pLUX in BV2 WT and *Smurf2* KO cells, transduced with EV or the *Smurf2* constructs shown in (A). Luminescence was measured at different timepoints (mean ± SD of a representative experiment performed in sextuplicate. ns, non-significant, *p ≤ 0.05 and **p ≤ 0.01; two-way ANOVA, with multiple comparisons).

To test the impact of SMURF2 complementation on Mtb replication, we infected the complemented cells with a luminescent Mtb strain encoding the entire firefly luciferase operon (Mtb-pLux) and quantified luminescence over three days. We used the Mtb-pLux strain in this and some subsequent experiments when testing multiple conditions simultaneously because the intracellular luminescence of the Mtb-pLux strain allows for rapid and continuous read-out of Mtb burden and comparable results to CFU plating (43, 44). As expected from prior CFU data (**Figures 1C, 3B**), BV2 *Smurf2* KO cells transduced with an empty vector exhibited reduced Mtb replication relative to BV2 wild type cells. This phenotype was reversed by complementation with *Smurf2^WT^* but not by *Smurf2^C716A^* (**Figure 4D**). While we cannot exclude the possibility that the inability of SMURF2^C716A^ to fully restore wild type phenotypes reflects reduced protein abundance rather than loss of catalytic function, these findings suggest that the E3 ligase activity of SMURF2 is required to promote intracellular Mtb replication in macrophages.

### SMURF2^WT^, but not SMURF2^C716A^, restores K48-linked ubiquitin colocalization with Mtb and LC3B recruitment to Mtb-containing structures in BV2 *Smurf2* KO macrophages

To determine whether SMURF2 regulates the recruitment of ubiquitin and autophagy machinery to Mtb, we assessed K48-linked ubiquitin and LC3B colocalization with Mtb-associated structures in BV2 *Smurf2* KO cells complemented with either *Smurf2^WT^* or *Smurf2^C716A^*. BV2 *Smurf2* KO cells transduced with an empty vector exhibited increased K48-linked ubiquitin colocalization with Mtb-associated structures **(Figures 5A, 5B)**, consistent with earlier observations **(Figure 2A)**. This phenotype was reversed by complementation with *Smurf2^WT^*, which restored K48-linked ubiquitin signal to levels observed in BV2 WT cells. In contrast, complementation with the catalytically inactive *Smurf2^C716A^* mutant failed to reduce K48 ubiquitin colocalization, indicating that SMURF2 ligase activity is required for this function **(Figures 5A, 5B)**.

**Figure 5:**
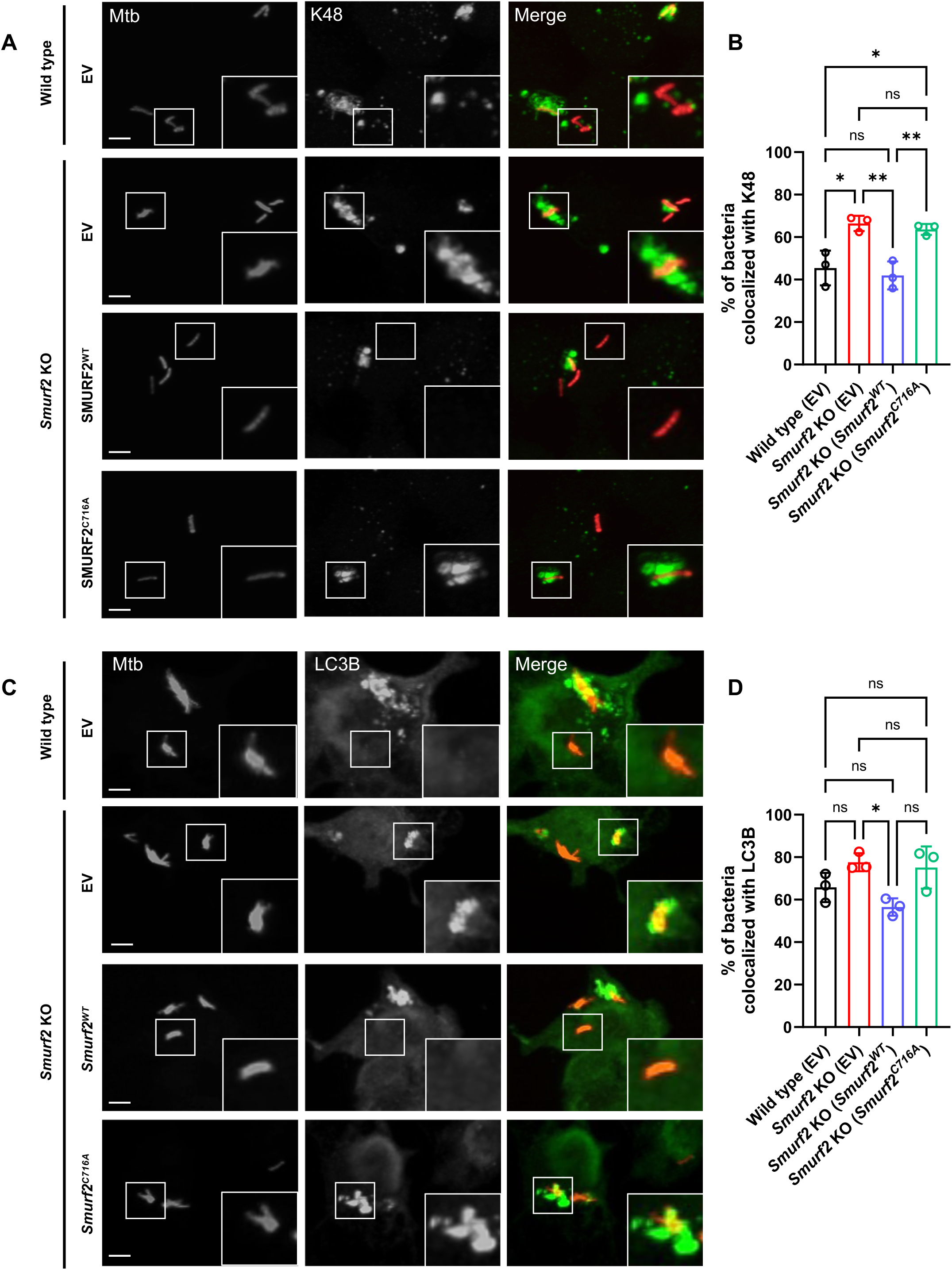
*Smurf2* complementation reduces K48 ubiquitin and LC3B colocalization with Mtb in murine macrophages. A) Representative immunofluorescence images of Mtb-mCherry (red) with K48 ubiquitin (green) in BV2 WT and *Smurf2* KO cells, transduced with either an empty vector (EV), or the constructs shown in Figure 4. Cells were stained with an anti-K48 antibody 17 h after infection. B) Quantification of Mtb-mCherry colocalization with K48 as shown in (A) (mean ± SD of a 3 independent experiments, performed in triplicate, ns, non-significant, *p ≤ 0.05 and **p ≤ 0.01; unpaired t-test). C) Representative immunofluorescence images of mCherry-expressing Mtb (red) with LC3B (green) under the same conditions as in (A), using an anti-LC3B antibody. D) Quantification of Mtb-mCherry colocalization with LC3B as shown in (C) (mean ± SD of a 3 independent experiments, performed in triplicate, ns, non-significant; *p ≤ 0.05; unpaired t-test). Scale bar, 5 µm.

We next examined LC3B recruitment to Mtb-associated structures. A modest increase in LC3B colocalization was again observed in BV2 *Smurf2* KO cells relative to wild type, although the difference did not reach statistical significance (**Figure 5C, 5D**). Complementation with *Smurf2^WT^* significantly reduced LC3B colocalization, restoring levels to those observed in BV2 wild type (EV) cells. In contrast, complementation with the *Smurf2^C716A^* mutant failed to reduce LC3B colocalization.

These results demonstrate that SMURF2 regulates the accumulation of K48-linked ubiquitin at Mtb-associated structures in a manner that depends on its catalytic activity, although differences in protein expression between SMURF2^WT^ and SMURF2^C716A^ may also contribute.

### Primary mouse bone marrow derived macrophages lacking *Smurf2* exhibit increased resistance to Mtb infection

To investigate the role of SMURF2 in primary macrophages, we generated mice with myeloid-specific deletion of *Smurf2* by crossing *Smurf2^flox/flox^* mice (45) with *LysMCre* transgenic mice (46). Western blot analysis of bone marrow–derived macrophages (BMDMs) confirmed recombination at the *Smurf2* locus, with an average 84% reduction in SMURF2 protein levels in *LysMCre*^+/-^ *Smurf2*^flox/flox^ BMDMs compared to littermate controls (**Supplemental Figure 1**).

To assess functional consequences of SMURF2 depletion, we infected *Smurf2^flox/flox^* and *LysMCre*^+/-^*Smurf2^flox/flox^* BMDMs with Mtb and quantified bacterial burden over a three-day time course. Like BV2 *Smurf2* KO cells, *LysMCre*^+/-^*Smurf2^flox/flox^* BMDMs showed reduced intracellular Mtb replication compared to controls at day three post-infection **(Figure 6A)**. We next examined additional molecular consequences of SMURF2 depletion. Immunoblot analysis confirmed reduced SMURF2 expression in *LysMCre*^+/-^ *Smurf2^flox/flox^* BMDMs following infection (**Figures 6B, 6C**). However, in contrast to BV2 *Smurf2* KO cells, while SMURF1 protein levels were modestly increased in *LysMCre*^+/-^ *Smurf2^flox/flox^* BMDMs compared to controls after Mtb infection, these results were not statistically significant (**Figures 6B, 6D**). LC3B lipidation was also not significantly different between genotypes (**Figures 6B, 6E**).

**Figure 6:**
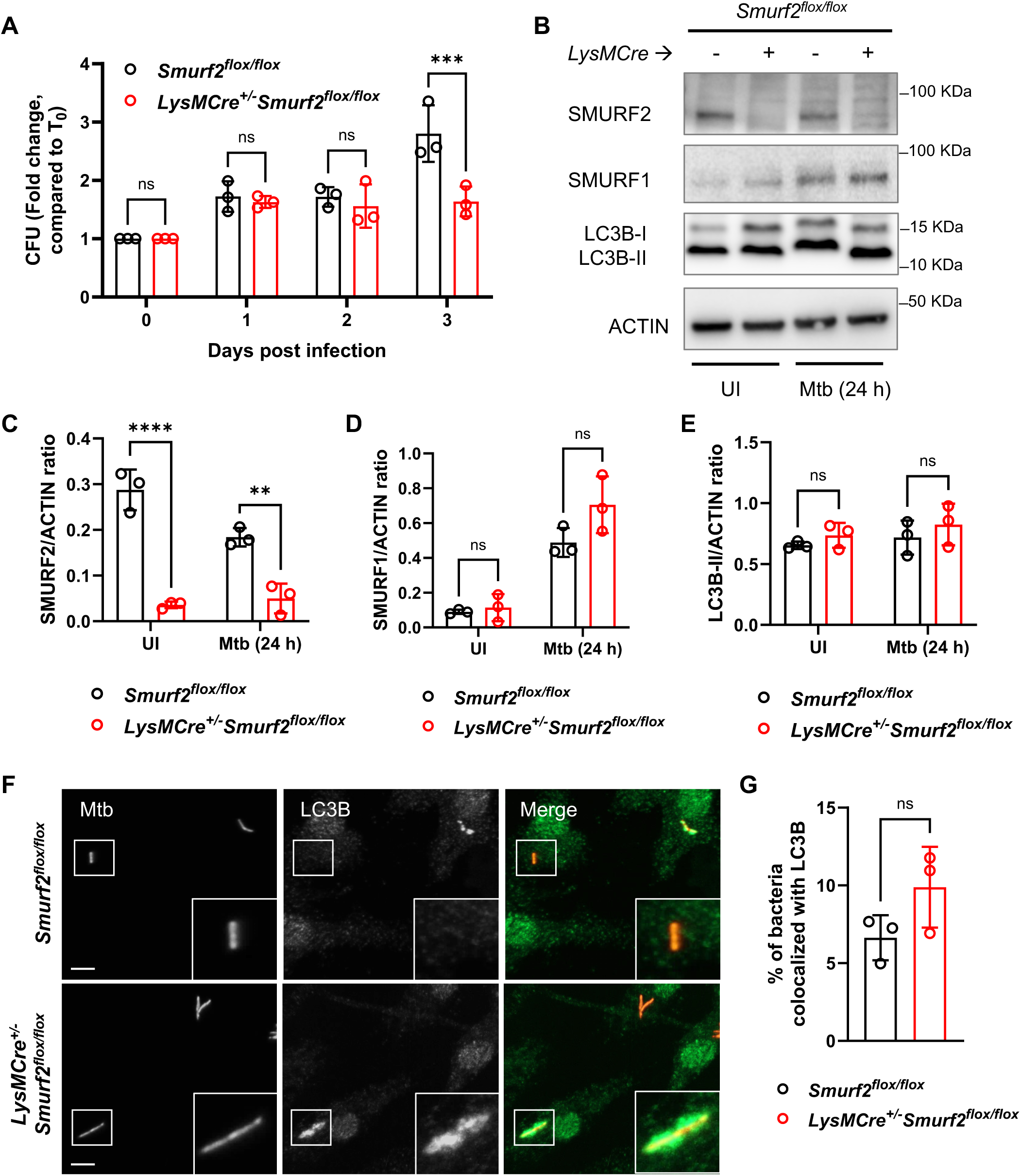
*LysMCre^+/-^Smurf2^flox/flox^* murine macrophages are more resistant to Mtb infection. A) Colony-forming units (CFU) of Mtb in *Smurf2^flox/flox^* and *LysMCre^+/-^ Smurf2^flox/flox^* BMDMs at different timepoints post-infection (ns, non-significant, and ***p < 0.001; two-way ANOVA). B) Representative immunoblot analysis showing SMURF2, SMURF1, and LC3B lipidation levels in protein lysates from *Smurf2^flox/flox^* and *LysMCre^+/-^ Smurf2^flox/flox^* BMDMs, 24 hours post infection. C-E) Densitometric quantification of SMURF2, SMURF1, and LC3B-II levels as shown in (B), from three independent experiments (mean ± SD of 3 independent experiments, ns, non-significant, **p < 0.01 and ****p < 0.0001; two-way ANOVA). F) Representative immunofluorescence images of Mtb-mCherry (red) with LC3B (green) in *Smurf2^flox/flox^* and *LysMCre^+/-^Smurf2^flox/flox^* BMDMs, using anti-LC3B antibody 17 hours after infection. G) Quantification of Mtb-mCherry colocalization with LC3B as shown in (F) (mean ± SD of 3 independent experiments, performed in triplicate, ns, non-significant; unpaired t-test). Scale bar, 5 µm.

To further assess autophagic targeting of Mtb, we performed immunofluorescence analysis of LC3B colocalization with Mtb-mCherry. As in BV2 cells, *LysMCre*^+/-^ *Smurf2^flox/flox^* BMDMs exhibited a modest increase in LC3B recruitment to Mtb-associated structures, but the difference was not statistically significant (**Figure 6F, 6G**).

These results indicate that partial depletion of SMURF2 in primary BMDMs is sufficient to reduce Mtb replication, although the associated changes in SMURF1 expression and LC3B lipidation are less pronounced than in BV2 *Smurf2* KO cells. Residual SMURF2 expression in *LysMCre^+/-^Smurf2^flox/flox^* BMDMs compared to BV2 *Smurf2* KO cells may contribute to the more modest phenotype.

### *Smurf2* does not impact bacterial burden or lung pathology during Mtb infection *in vivo*

We next tested whether *Smurf2* contributes to host defense against Mtb infection *in vivo*. *LysMCre*^+/-^*Smurf2^flox/flox^* and littermate *Smurf2^flox/flox^* control mice were infected via aerosol with a low-dose inoculum of Mtb (∼50 CFU) (**Figure 7A**) (47–49). At 7, 21, and 42 days post-infection, we quantified bacterial burden in the lungs, spleen, and liver. No significant differences in CFU were observed at any time point between groups **(Figures 7B-D)**. To assess the impact of *Smurf2* deletion on tissue pathology, lung sections were examined by histology at days 21 and 42 post-infection. *LysMCre*^+/-^*Smurf2^flox/flox^* mice showed similar degrees of inflammation compared to *Smurf2^flox/flox^* controls **(Figures 7E-G)**.

**Figure 7:**
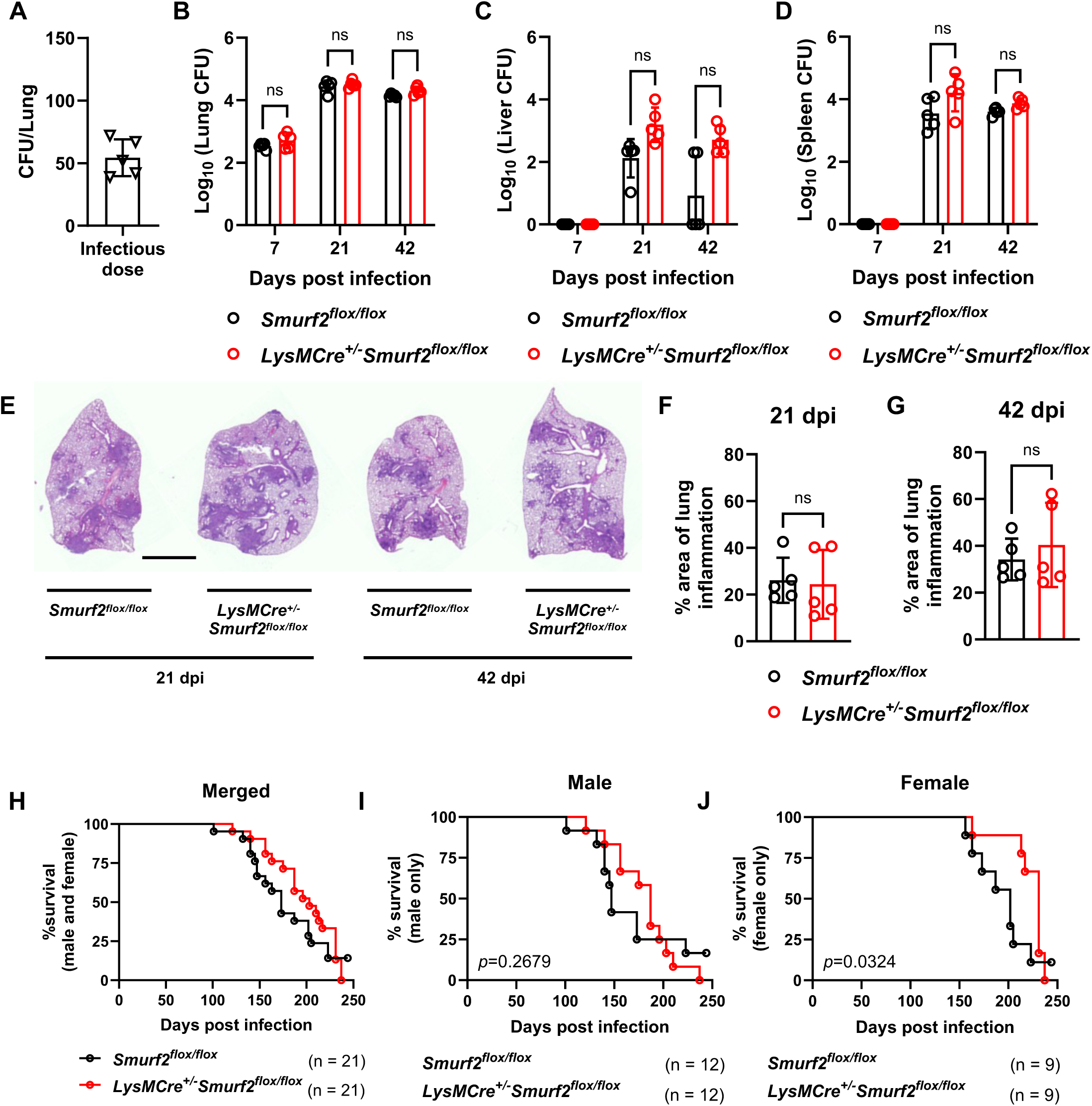
*Smurf2* deficiency does not significantly impact Mtb infection *in vivo*. A) *Smurf2^flox/flox^* and *LysMCre^+/-^Smurf2^flox/flox^* mice were aerosol-infected with ∼50 CFU Erdman Mtb. Mice were euthanized at different timepoints, and CFUs were enumerated in the (B) lungs, (C) liver, and (D) spleen at 7, 21, and 42 days post-infection (dpi) (n = 5 mice/group, mean ± SD of an experiment performed in quadruplicate, ns. p > 0.05; two-way ANOVA). E) Representative hematoxylin and eosin staining of lung sections from Mtb-infected mice at 21 and 42 dpi. Scale bar, 2.5 mm. F, G) Quantification of inflammatory areas in lung sections at 21- and 42-dpi, respectively, as shown in (E). Data represent mean ± SD (n = 5 mice/group, ns. p > 0.05, Mann-Whitney test). H-J) Kaplan-Meier survival curves of *Smurf2^flox/flox^* and *LysMCre^+/-^Smurf2^flox/flox^* Mtb-infected mice (n = 21 animals/genotype), shown for (H) male and female mice, (I) male only, and (J) female only (exact p values are shown on the graph; Log-rank test).

We then monitored survival following aerosol infection. When male and female mice were analyzed together, survival curves were not significantly different (**Figure 7H**). However, sex-stratified analysis revealed that female *LysMCre*^+/-^*Smurf2^flox/flox^* mice survived significantly longer than female *Smurf2^flox/flox^* littermates, while no survival difference was observed among males **(Figures 7I, J).**

These data indicate that myeloid-specific deletion of *Smurf2* via *LysMCre* expression does not alter bacterial burden or lung pathology during acute Mtb infection but may modestly improve survival in a sex-dependent manner.

### *SMURF2* knockdown reduces Mtb intracellular replication in primary human macrophages

To determine whether SMURF2 contributes to Mtb control in human macrophages, we differentiated monocyte-derived macrophages (hMDMs) from peripheral blood CD14^+^-monocytes isolated from healthy donors. hMDMs were transduced with lentiviral vectors encoding a non-targeting control short hairpin RNA (shRNA) or one of two independent shRNAs targeting *SMURF2* **(Figure 8A)**. Knockdown efficiency was analyzed by immunoblot, with varying degrees of SMURF2 depletion observed across the two shRNA constructs **(Figure 8B)**. We then infected hMDMs with Mtb and quantified intracellular bacterial burden over a three-day time course by CFU analysis. Both *SMURF2*-targeting shRNAs significantly reduced Mtb replication compared to the non-targeting control (**Figure 8C**). Notably, cells transduced to express the shRNA that achieved more efficient SMURF2 knockdown (shRNA#2) also demonstrated a lower bacterial burden, suggesting a dose-dependent relationship between SMURF2 expression and Mtb growth in hMDMs. These data demonstrate that SMURF2 negatively regulates macrophage control of Mtb in primary human cells, supporting a conserved role for SMURF2 in suppressing cell-autonomous immunity across species.

**Figure 8:**
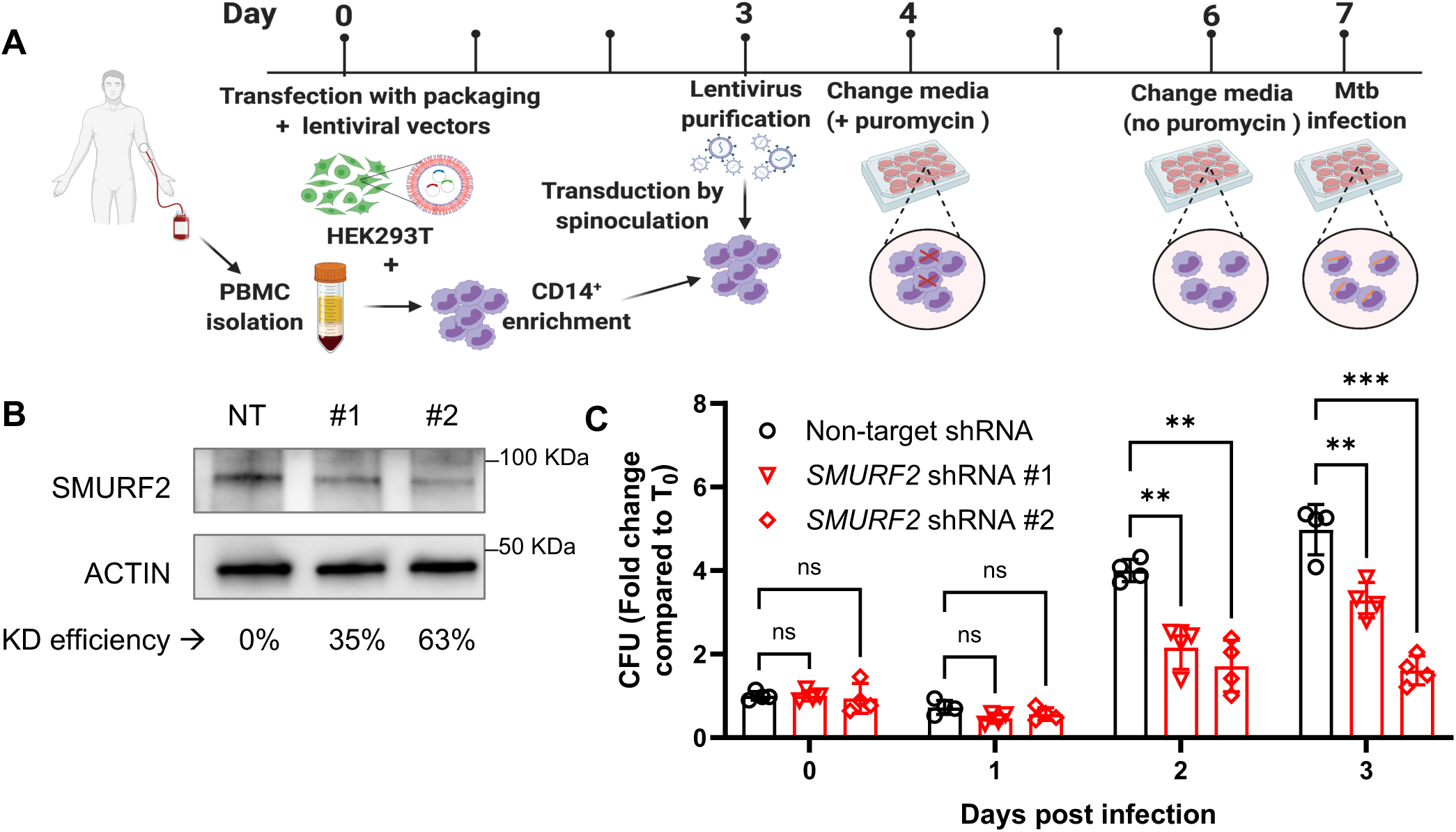
*SMURF2* knockdown reduces Mtb intracellular growth in primary CD14-positive human macrophages. A) Schematic of the isolation and transduction of CD-14^+^-enriched primary human macrophages (see Material and Methods for details). B) Representative immunoblot of SMURF2 in hMDMs from a single donor transduced with non-targeting shRNA (NT) or two different *SMURF2* shRNA (#1 or #2). SMURF2 knockdown efficiencies are indicated below ACTIN bands. C) Intracellular Mtb CFU in hMDMs transduced as in (A), at different timepoints post-infection (mean ± SD of a representative experiment performed in quadruplicate, ns. p > 0.05, **p ≤ 0.01 and ***p ≤ 0.001; two-way ANOVA).

### Pharmacologic inhibition of SMURF2 reduces Mtb replication and enhances LC3B recruitment in human macrophages

To evaluate whether pharmacologic inhibition of SMURF2 could enhance host control of Mtb, we treated hMDMs with Heclin, a small molecule that targets multiple HECT-type E3 ligases including SMURF2 (50). We infected hMDMs with Mtb-pLux, treated cells with increasing concentrations of Heclin, and measured luminescence over three days. Heclin reduced intracellular Mtb replication in a dose-dependent manner at all time points **(Figure 9A)**. To determine whether Heclin directly affects Mtb viability, axenic Mtb cultures were incubated with Heclin or DMSO, and growth was measured by optical density. Heclin alone had no impact on axenic Mtb growth, confirming that the reduction in luminescence observed in macrophages was not due to a direct antimicrobial effect (**Supplemental Figure 2**).

**Figure 9:**
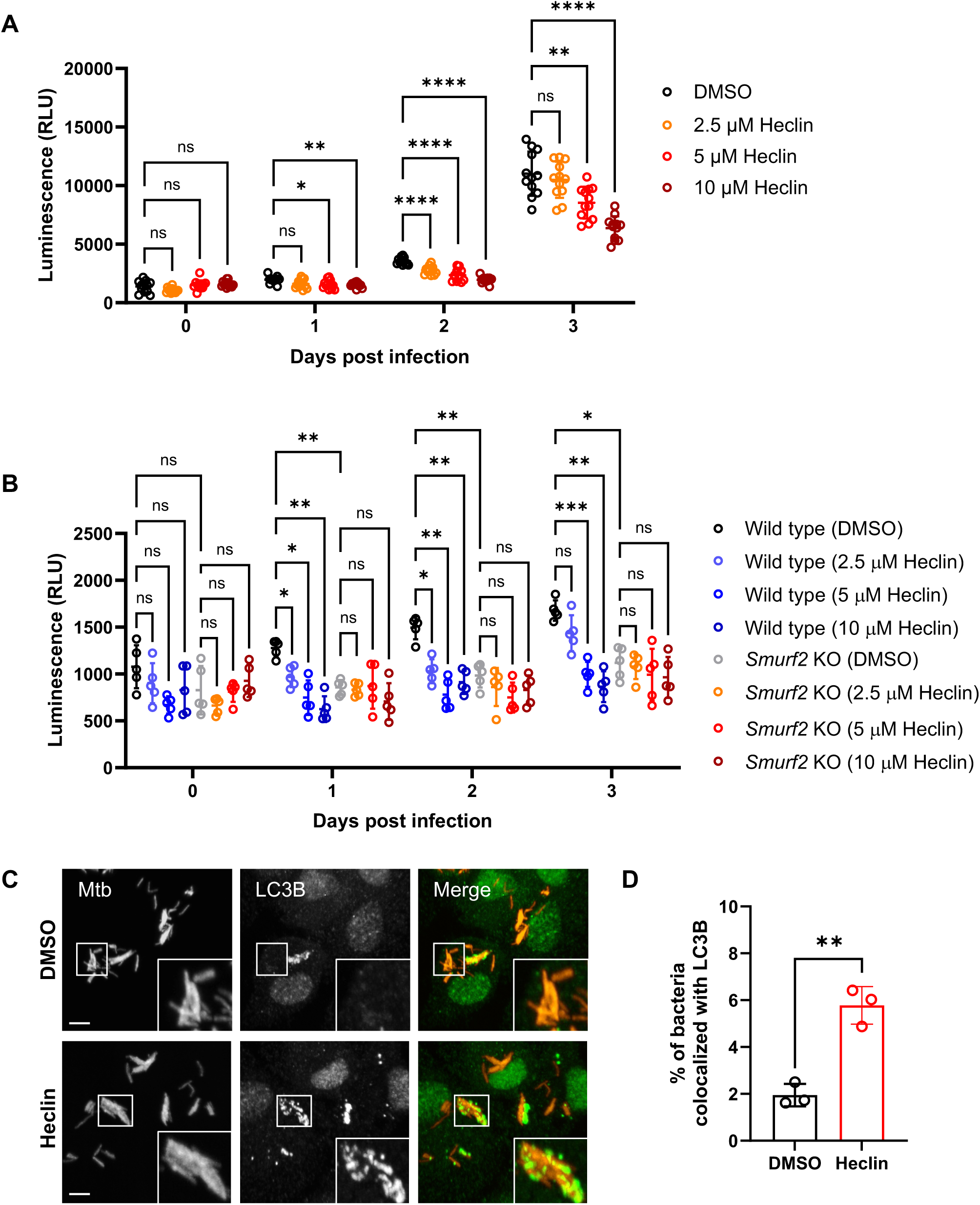
Inhibition of SMURF2 by Heclin reduces Mtb intracellular growth in primary CD14^+^ human macrophages. A) Luminescence from Mtb-pLUX in hMDMs exposed to either 0.1% DMSO (vehicle control) or increasing concentrations of Heclin, as indicated (see Material and Methods for details). Luminescence was measured at different timepoints (mean ± SD of a representative experiment performed in quadruplicate, ns. p > 0.05, *p ≤ 0.05, **p ≤ 0.01 and ****p ≤ 0.0001; two-way ANOVA). B) Luminescence from Mtb-pLUX in BV2 WT and *Smurf2* KO cells exposed to either 0.1% DMSO (vehicle control) or increasing concentrations of Heclin. Luminescence was measured at different timepoints (mean ± SD of a representative experiment performed in quintuplicate. ns. p > 0.05, *p ≤ 0.05, **p ≤ 0.01 and ***p ≤ 0.001; two-way ANOVA, with multiple comparisons). C) Representative immunofluorescence images of Mtb-mCherry (red) and LC3B (green) in human macrophages exposed to either 0.1% DMSO (vehicle control) or 10 µM Heclin. D) Quantification of Mtb-mCherry colocalization with LC3B shown in (C) mean ± SD of a 3 independent experiments, performed in triplicate, **p ≤ 0.01; unpaired t-test). Scale bar, 5 µm.

Since Heclin inhibits several HECT ligases, including SMURF2 (IC₅₀ 6.8 µM), NEDD4 (IC₅₀ 6.3 µM), and WWP1 (IC₅₀ 6.9 µM) (50), we sought to determine whether its antibacterial effect in macrophages was mediated specifically through SMURF2. To do this, we infected both BV2 wild-type and BV2 *Smurf2* KO cells with Mtb-pLux and treated them with increasing concentrations of Heclin. Like hMDMs, BV2 wild-type cells exhibited a dose-dependent reduction in Mtb replication, whereas BV2 *Smurf2* KO cells were unresponsive to Heclin treatment but showed reduced bacterial levels similar to the highest tested Heclin concentration (**Figure 9B**). These results suggest that SMURF2 is the relevant HECT E3 ligase mediating the effect of Heclin in restricting Mtb replication.

To assess the effect of SMURF2 inhibition on autophagy, we quantified LC3B colocalization with Mtb-associated structures in Heclin- or vehicle-treated hMDMs. Heclin treatment significantly increased LC3B recruitment to Mtb-mCherry in hMDMs, consistent with enhanced selective autophagy **(Figures 9C, D)**. We also attempted to assess K48-linked ubiquitin colocalization, but the K48-ubiquitin staining was inconsistent and could not be reliably quantified.

Together, these findings support a role for SMURF2 as a negative regulator of selective autophagy in human macrophages and demonstrate that its pharmacologic inhibition enhances both murine and human macrophage control of Mtb.

## Discussion

Our findings identify SMURF2 as a negative regulator of autophagy-mediated, cell-autonomous immunity to Mtb. In murine macrophages, *Smurf2* deletion increased SMURF1 abundance, enhanced K48-linked ubiquitination and reduced bacterial replication. We also observed a trend towards increased LC3B recruitment to Mtb associated structures in murine macrophages lacking SMURF2. These effects were lost in the absence of SMURF1, consistent with prior evidence that SMURF2 promotes SMURF1 degradation (23) and indicating that SMURF2 limits autophagy primarily by destabilizing SMURF1. We extended these observations to primary human macrophages, where both genetic and pharmacologic inhibition of SMURF2 increased LC3B recruitment and reduced Mtb growth without direct antibacterial effects on axenic growth. The concordance between murine and human data highlights a conserved mechanism across species and underscores the potential of SMURF2 targeting as a host-directed therapeutic approach for tuberculosis.

E3 ubiquitin ligases, including SMURF2, are central regulators of protein ubiquitination, influencing diverse immune processes such as lymphocyte development, antigen presentation, and immune evasion (51–53). While their antiviral functions are well described (27, 54, 55), the role of HECT E3 ligases in antibacterial immunity is less understood. Several ligases, including TRIM32, NEDD4, and SMURF1, promote autophagy-mediated Mtb clearance by enhancing bacterial ubiquitination or stabilizing key autophagy proteins such as BECN1 (18, 56). Other ligases, such as TRIM27, promote bacterial clearance by activating transcription factors like TFEB, independent of their ligase activity (57).

Mtb can also subvert host E3 ligases to favor its survival. For example, the Mtb PE/PPE protein PE5 hijacks the CRL2 complex to suppress cell-autonomous immunity and promote its intracellular replication (58), while Mtb Rv0222 interacts with ANAPC2 to dampen proinflammatory cytokine production (59). Finally, PPE36 promotes SMURF1-dependent ubiquitination and proteasomal degradation of MyD88, limiting host inflammatory responses (60).

While SMURF2 deficiency enhanced bacterial clearance *in vitro*, its *in vivo* effects were more modest: overall bacterial burden and lung inflammation were unchanged, but female *LysMCre^+/-^Smurf2^flox/flox^* mice exhibited prolonged survival compared to littermate controls. This observation, together with known sex-based differences in TB susceptibility (49, 61–63) and autophagy regulation (64), suggests that SMURF2 function may be influenced by biological sex and warrants further study.

Our experiments in human macrophages underscore the translational importance of these findings. SMURF2 knockdown or pharmacologic inhibition with Heclin significantly reduced Mtb replication without direct antibacterial effects, and increased LC3B colocalization with Mtb. The absence of the Heclin effect in *Smurf2*-deficient cells supports SMURF2 as its relevant target in this context, highlighting the potential for SMURF2 inhibitors as host-directed therapeutics.

Although autophagy has been widely studied as a cell-autonomous defense against intracellular pathogens, including Mtb, there is a lack of consensus from *in vivo* models regarding its protective function (65–68). Conflicting results from murine models may reflect differences in Mtb strain used, infectious dose, mouse background, specificity of gene deletion and duration of survival studies. Moreover, Mtb encodes virulence factors that modulate autophagy or promote autophagy evasion. For example, the secreted effector <u>e</u>nhanced intracellular <u>s</u>urvival (Eis) inhibits autophagy initiation through modulation of host redox-dependent signaling and acetylation pathways (69, 70), while PE_PGRS47 inhibits autophagy initiation by interacting with Ras-related protein Rab1a (71, 72). Likewise, the SecA2 dependent protein export system exports effectors including protein kinase G (PknG) and secreted acid phosphatase (SapM) to block phagosome-lysosome fusion through blockade of the small GTPase Rab7 (73, 74). Mtb lipids, including phthiocerol dimycocerosates (PDIM) and sulfolipid-1 (SL-1), further impair phagosome maturation and autophagic targeting (75–78). These strategies underscore the importance of autophagy as a barrier to infection and suggest that Mtb has evolved multiple mechanisms to subvert it.

In contrast to the debated role of autophagy in TB pathogenesis in mice, evidence from human macrophages consistently supports its contribution to cell-autonomous defense. Genetic ablation of *SMURF1* (*10*), *IRGM* (*79*), *ATG7* (80) or *ATG14* (80) in human macrophages increases mycobacterial replication, whereas inhibition of the deubiquitinase USP15, which counters PARKIN, decreases Mtb replication (81). Our findings add to this body of evidence by showing that both genetic and pharmacologic disruption of SMURF2 enhances Mtb clearance in primary human macrophages by stimulating autophagy. In addition, recent work demonstrates that calcium leakage following Mtb phagosome damage induces multimeric ATG8/LC3 lipidation, limiting bacterial replication through a noncanonical autophagy process (82). Together, these findings underscore that selective autophagy, through both canonical and noncanonical mechanisms, is a key component of human host defense against Mtb and a promising target for therapeutic intervention.

In summary, our data define SMURF2 as a negative regulator of autophagy-mediated control of Mtb and provide mechanistic insight into how E3 ligases shape cell-autonomous immunity. These findings support further exploration of SMURF2 as a therapeutic target for TB, particularly in combination with standard antimicrobial regimens.

## Material and Methods

### Mice

All animal experiments were approved by the Institutional Animal Care and Use Committee (IACUC) at the University of Texas Southwestern Medical Center (UTSW) and were conducted in accordance with the NIH Guide for the Care and Use of Laboratory Animals (8^th^ edition, National Academies Press, 2011). UTSW is accredited by the American Association for Accreditation of Laboratory Animal Care (AAALAC). To obtain *LysMCre^+/-^Smurf2^flox/flox^* mice, we crossed *LysMCre mice* (strain 004781, Jackson Laboratory, Bar Harbor, ME) (46) with *Smurf2^flox/flox^* mice (a gift from Ying E. Zhang, National Cancer Institute, National Institutes of Health, Bethesda, MD (45)). Genotyping of mice was performed according to the protocols provided for each strain on The Jackson Laboratory website. Briefly, genomic DNA was extracted from ear punches using a previously described protocol (83). All mice were maintained in a pathogen-free facility with a 12-hour light/dark cycle and fed *ad libitum* in accordance with institutional guidelines.

### Bacterial strains

We used the Erdman strain of *Mycobacterium tuberculosis* (Mtb) as the parental strain for this study. The Mtb mCherry-expressing strain has been described previously (10, 47, 84). The Erdman luminescent reporter strain was generated by transforming the parental Mtb strain with pMV306hsp+LuxG13 plasmid (a gift from Brian Robertson & Siouxsie Wiles (Addgene plasmid # 26161; http://n2t.net/addgene:26161; RRID:Addgene_26161) (85). Mtb cultures were grown to mid-log phase in Middlebrook 7H9 medium (Difco), supplemented with 10% Middlebrook OADC (BD), 0.5% glycerol, and 0.05% Tween-80 at 37°C using aseptic techniques in a Biosafety Level 3 laboratory. Strains were propagated with minimal passage to maintain virulence. Following growth of Mtb, bacteria were washed with phosphate-buffered saline (PBS) and sonicated with three pulses lasting 7 seconds each, with an amplitude of 90% and 5-second rests between pulses. After sonication, the concentration of bacterial suspensions was determined by measuring optical density at 600 nm (OD_600_), using the formula: 1 OD_600_ = 3 x 10^8^ bacteria/mL, and diluted in macrophage media prior to infection.

To assess the effect of the HECT E3 ligase inhibitor Heclin on Mtb growth, Erdman strain cultures were grown in 7H9 medium supplemented with 10% OADC, 0.5% glycerol, and 0.05% Tween-80 in the presence of either 10 µM Heclin or 0.1% DMSO (vehicle control). Bacterial growth was monitored by measuring OD_600_ at different time points.

### Cell culture

The mouse microglial cell line BV2 was cultured in complete Dulbecco’s modified Eagle medium (DMEM) supplemented with 10% fetal bovine serum (FBS) (Gibco), and 10 mM HEPES (Thermo Fisher) and maintained in 5% CO_2_ at 37°C. Bone marrow-derived macrophages (BMDMs) were isolated and cultured from 8-12 week old mice femurs and tibia as previously described (47). Peripheral blood mononuclear cells (PBMCs) were isolated from buffy coats of healthy donors using SepMate^TM^ tubes (StemCell Technologies, Vancouver, Canada) and enriched for CD14⁺ monocytes using MACS CD14⁺ microbeads (Miltenyi), following the manufacturer’s protocol. Adherent human macrophages (hMDMs) were obtained by culturing CD14^+^ monocytes in complete RPMI 1640 (supplemented with 10% FBS, 2 mM L-glutamine, 100 U/mL penicillin, 100 μg/mL streptomycin), with 10% inactivated human serum, and 50 ng/mL recombinant human M-CSF (R&D Systems) for 7 days. Lentiviral transductions for *SMURF2* knockdown were performed using the protocol described below.

### Lentiviral transduction for Smurf2 overexpression in BV2 cells

To achieve transgenic expression of SMURF2 in BV2 cells, we cloned full-length *Smurf2* cDNA (*Smurf2^WT^* or catalytic-dead *Smurf2^C716A^*) (86) into the entry vector pENTR/D-TOPO, which contains a Myc-tag sequence. These constructs were then subcloned into the lentiviral GATEWAY destination vector pLENTI CMV Blast using the LR Clonase™ II enzyme mix (Thermo Fisher). All plasmids were sequenced to verify correct gene insertion. Lentiviral particles were generated by transfecting HEK293T cells with the lentiviral constructs along with the packaging vectors pMD2.G and psPAX2 using FuGene HD transfection reagent (Promega), following the manufacturer’s protocol. After 48 hours of incubation in 5% CO₂ at 37°C, virus-containing supernatants were collected, filtered through a 0.45 µm filter (Millipore), and added to BV2 cell cultures in the presence of 8 µg/mL Polybrene (Millipore). Cells were incubated in 5% CO₂ at 37°C for 16 hours, when the medium was replaced with fresh complete DMEM containing 4 µg/mL blasticidin. The medium was changed every other day until resistant cell populations were established.

### Lentiviral Transduction for SMURF2 Knockdown in human monocyte-derived macrophages

shRNAs were used to induce sequence-specific *SMURF2* knockdown in hMDMs. shRNA synthetic molecules (clone TRCN0000003477, target sequence: CGCCTCAAAGACACTGGTTAT; clone TRCN0000003478, target sequence: CCACCCTATGAAAGCTATGAA) were obtained from Sigma, with a non-targeting sequence serving as a control. Lentiviral particles containing shRNAs were produced in HEK293T cells as described previously.

Human MDMs were transduced on day 4 of differentiation by adding lentivirus-containing supernatants to the cultures in the presence of 8 µg/mL Polybrene (Millipore). Cells were transduced via spinoculation at 800 × g for 1 hour at 30°C. Following transduction, the medium was replaced with fresh complete RPMI 1640 supplemented with 50 ng GM-CSF. On day 5, the medium was replaced with GM-CSF-free RPMI 1640 containing 2 µg/mL puromycin for selection. On day 7, MDMs were infected with Mtb, as described below. Knockdown (KD) efficiency was calculated as the percentage decrease in the SMURF2/ACTIN ratio in *SMURF2* shRNA-transduced cells relative to Non-target shRNA-transduced cells.

### Intracellular quantification of Mtb

To enumerate the number of intracellular bacteria by colony-forming units (CFU) method, we infected macrophages with Mtb at a multiplicity of infection (MOI) of 1. After phagocytosis, cells were washed twice with PBS and lysed at different time points with 0.5% Triton X-100 in PBS. Serial dilutions were plated on 7H11 plates supplemented with 10% OADC, 0.5% glycerol, and 50 µg/mL cycloheximide, and grown for 3 to 4 weeks at 37°C.

To monitor intracellular mycobacterial growth by luminescence, we infected macrophages with the luminescent reporter strain at a MOI of 3, and after phagocytosis, cells were washed twice with PBS and fresh media was added to each well prior to the measurement. Luminescence was measured using a BioTek Synergy Neo2 Hybrid Multimode Reader every 24 hours for 3 days.

To assess the effect of the HECT E3 ligase inhibitor Heclin (Sigma) on intracellular Mtb quantification, cells were treated with different concentrations of the inhibitor (diluted in DMSO) after phagocytosis and before cell lysis for Western blot or bacterial growth in macrophages.

### Western blot

Cells were seeded at 5 x 10^5^ cells per well in 12-well plates and maintained at 37°C with 5% CO_2_ for 4 hours prior to infection with parental Mtb at a MOI of 5. After phagocytosis, cells were maintained in complete DMEM with or without 0.1% DMSO or 10 µM Heclin. At different time points, cells were washed twice with PBS and lysed in RIPA buffer (Cell Signaling) supplemented with cOmplete EDTA-free protease inhibitor cocktail (Roche) and Halt^TM^ phosphatase inhibitor cocktail (Thermo Fisher Scientific). Lysates were incubated on ice for 20 minutes, then collected and filtered twice through a 0.2 µm filter (Millipore). Laemmli sample buffer was added, and samples were boiled for 3 minutes.

Proteins were separated by SDS-PAGE using 4-20% polyacrylamide resolving gel (Bio-Rad). After electrophoresis, proteins were transferred to PVDF membranes and blocked with 5% BSA in Tris-buffered saline with 0.1% Tween-20 (TBS-T). To detect SMURF2, SMURF1, and LC3B, the following antibodies were used: anti-LC3B (Novus), SMURF2 (Cell Signaling), SMURF1 (Santa Cruz), respectively. Anti-β-ACTIN (Santa Cruz) was used as a loading control. Membranes were developed using Clarity Western ECL Substrate (Bio-Rad).

### Immunofluorescence analysis

Cells were seeded at 1 x 10^5^ cells per well in 24-well plates with glass coverslips and maintained at 37°C with 5% CO_2_ for 4 hours prior to infection with mCherry-expressing Mtb at a MOI of 1 for 17 hours. Coverslips were then washed twice with PBS, fixed in 1% paraformaldehyde for 16 hours at 4°C, and washed twice again in PBS. For K48 ubiquitin staining, coverslips were permeabilized with 0.3% Triton-X-100 in PBS (PBS-T) for 15 min, blocked with 3% BSA in PBS-T for 1 h and incubated with humanized anti-K48 ubiquitin antibody (Genentech) for 1 h at room temperature. Coverslips were then washed 3x with PBS and incubated with Alexa 488-conjugated goat anti-human IgG (Thermo Fisher) for 1 h at room temperature, protected from light. For LC3B staining, coverslips were permeabilized with 100% methanol for 6 min at −20°C, blocked with 3% BSA in PBS for 1 h and incubated with rabbit anti-LC3B (Novus) for 1 h at room temperature. Coverslips were then washed 3x with PBS and incubated with Alexa 488-conjugated goat anti-rabbit IgG (Invitrogen) for 1 h at room temperature, protected from light. After three additional washes with PBS, all coverslips were mounted using ProLong^TM^ Diamond Antifade Mountant with DAPI and sealed with nail polish. Slides were visualized using a Nikon CSU-W1 SoRa spinning disk confocal microscope with a 60x objective. Colocalization event counting was performed in a blind manner, with samples numbered to prevent investigator bias.

### Low dose aerosol infection of mice with M. tuberculosis

*Smurf2^flox/flox^* and *LysMCre^+/-^Smurf2^flox/flox^* mice were infected via aerosol as previously described (47, 65). Briefly, mid-log-phase parental Mtb were washed three times with PBS, sonicated as described above and resuspended in PBS to an OD_600_ of 0.1 in PBS. Bacterial suspensions were used to infect mice in a GlasCol aerosol chamber (GlasCol Inc.), with an inoculum of approximately 50 bacteria per mouse. On day 0, both lungs from five mice were plated to determine the initial bacterial load. At 7, 21, and 42 days post-infection, lungs, liver, and spleen were collected, homogenized in PBS, and plated on 7H11 agar plates for CFU enumeration. For survival studies, mice were weighed weekly and euthanized upon reaching 15% loss of their maximum body weight, threshold marker of imminent death, as has been previously reported (47).

### Histological analysis

The left lung lobe from each mouse was collected at indicated time points and fixed in 10% formaldehyde in phosphate-buffered saline (PBS), pH 7.2. Tissues were then transferred to 70% ethanol and submitted to the Molecular Pathology Core Facility at UT Southwestern for paraffin embedding, sectioning (5 μm thickness), and Hematoxylin & Eosin staining. Slides were scanned at 20X magnification using a NanoZoomer S60 digital slide scanner. The percentage of inflammation per tissue was subsequently calculated using ImageJ by observers blinded to mouse genotype.

### Statistical analysis

All statistical analysis was conducted using GraphPad Prism software (version 10). For *in vitro* studies, data were analyzed using unpaired, 2-tailed t-test for comparisons between two groups, and analysis of variance (ANOVA) for comparisons with more than two groups. For *in vivo* studies, the Mann-Whitney U test was used for nonparametric comparisons between 2 groups of mice. For mouse mortality studies, Kaplan-Meyer survival curves were generated and analyzed through log-rank tests to determine statistical significance. Densitometric analysis of immunoblots was performed using ImageJ software (version 1.54f). The specific statistical tests used are indicated in the figure legends.

## Acknowledgements

We thank Ying E. Zhang (NCI/CCR, Bethesda, MD, USA) for the *Smurf2^flox/flox^* mice, and Luis Franco (Federal University of Minas Gerais, Belo Horizonte, Brazil) for valuable feedback on experiments. The authors also thank core facilities at UT Southwestern Medical Center for their important contributions to this work. We thank the UT Southwestern Quantitative Light Microscopy Core, particularly the core director Marcel Mettlen, a Shared Resource of the Harold C. Simmons Cancer Center, supported in part by an NCI Cancer Center Support Grant, 1P30 CA142543-01. The Nikon SoRa spinning disk microscope was purchased with a shared instrumentation grant from NIH, 1S10OD028630-01 to Katherine Luby-Phelps. We also thank the Histo Pathology Core, particularly John Shelton and the Whole Brain Microscopy Facility core director Denise Ramirez.

## Author contributions

Conceptualization, P.C., M.U.S.; Formal analysis, P.C., M.U.S.; Funding acquisition, M.U.S.; Investigation, P.C., K.R., V.E., B.D., K.N., S.A.; Project administration, M.U.S.; Supervision, M.U.S.; Visualization, M.U.S.; Writing – original draft, P.C., M.U.S.; Writing – review and editing, All authors.

