## Supplemental Figures and Legends for "SMURF2 inhibits autophagic control of *Mycobacterium tuberculosis* in macrophages"

### Supplemental Figure 1

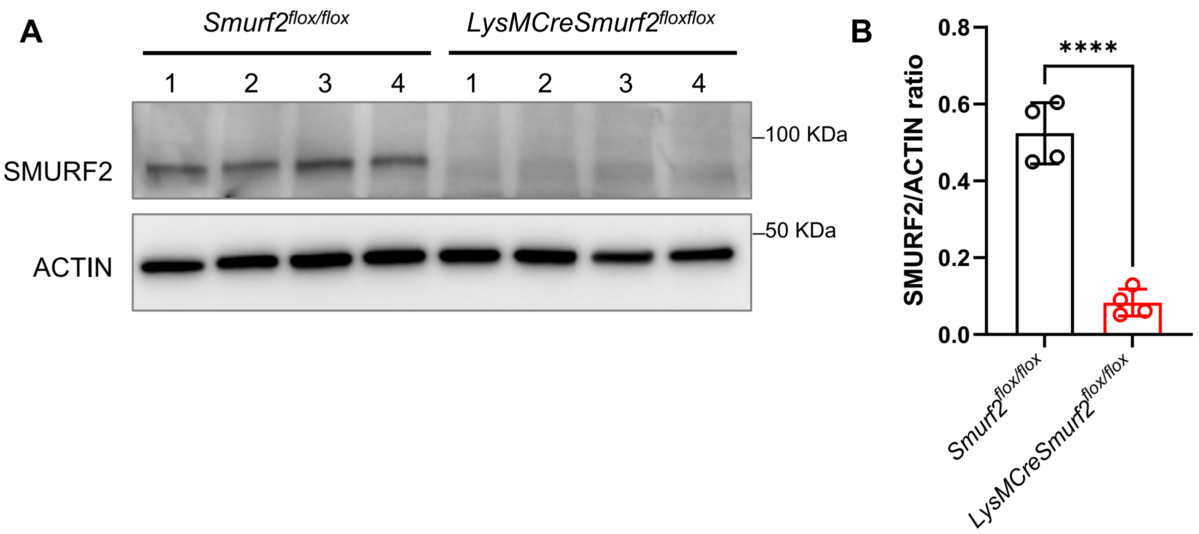

### Supplemental Figure 2

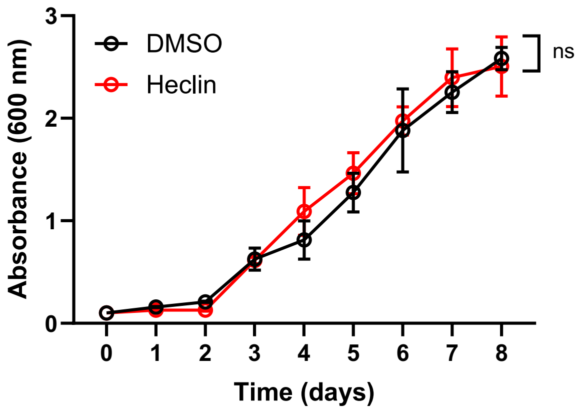

**Supplemental Figure 1: SMURF2 expression in *Smurf2*$^{flox/flox}$ and *LysMCre*$^{+/-}$ *Smurf2*$^{flox/flox}$ in murine bone-marrow-derived macrophages (BMDM).**

A) Representative immunoblot analysis of SMURF2 in protein lysates from *Smurf2*$^{flox/flox}$ and *LysMCre*$^{+/-}$*Smurf2*$^{flox/flox}$ BMDM from four mice per genotype, under basal conditions. B) Densitometric quantification of SMURF2 levels shown in (A) (n = 4 mice/group, ***p < 0.0001; unpaired t-test).

**Supplemental Figure 2: Mtb growth in the presence of DMSO or Heclin.**

Growth curves of Mtb Erdman in 7H9 medium, supplemented with 10% OADC, 0.5% glycerol, and 0.05% Tween-80, exposed to either 0.1% DMSO (vehicle control) or 10 µM Heclin. Absorbance at 600 nm was measured at different timepoints. Data represent mean ± SD from three independent cultures per condition (ns. p > 0.05; unpaired t-test).
